# 20-Hydroxyecdysone activates the protective arm of the renin angiotensin system via Mas receptor

**DOI:** 10.1101/2020.04.08.032607

**Authors:** René Lafont, Sophie Raynal, Maria Serova, Blaise Didry-Barca, Louis Guibout, Mathilde Latil, Pierre J. Dilda, Waly Dioh, Stanislas Veillet

## Abstract

20-Hydroxyecdysone (20E) is a steroid hormone that plays a key role in insect development through nuclear ecdysone receptors (EcRs) and at least one membrane GPCR receptor (DopEcR) and displays numerous pharmacological effects in mammals. However, its mechanism of action is still debated, involving either an unidentified GPCR or the estrogen ERβ receptor. The goal of our study was to better understand 20E mechanism of action.

A mouse myoblast cell line (C2C12) and the gene expression of myostatin (a negative regulator of muscle growth) was used as a reporter system of anabolic activity. Experiments using protein-bound 20E established the involvement of a membrane receptor. 20E-like effects were also observed with Angiotensin-(1-7), the endogenous ligand of Mas. Additionally, the effect on myostatin gene expression was abolished by Mas receptor knock-down using small interfering RNA (siRNA) or pharmacological inhibitors.

17β-Estradiol (E2) also inhibited myostatin gene expression, but protein-bound E2 was inactive, and E2 activity was not abolished by angiotensin-(1-7) antagonists. A mechanism involving cooperation between Mas receptor and a membrane-bound palmitoylated estrogen receptor is proposed.

The possibility to activate the Mas receptor with a safe steroid molecule is consistent with the pleiotropic pharmacological effects of ecdysteroids in mammals and indeed this mechanism may explain the close similarity between angiotensin-(1-7) and 20E effects. Our findings open a lot of possible therapeutic developments by stimulating the protective arm of the renin-angiotensin-aldosterone system (RAAS) with 20E.

## INTRODUCTION

### Steroids in animal and plant kingdoms

Steroid hormones are (chole)sterol derivatives widespread in animals and plants, where they are involved in the control of plenty of physiological processes. They include for example vertebrate sex hormones (progestagens, estrogens, androgens), insect moulting hormones (ecdysteroids), as well as plant growth hormones (brassinosteroids). This means that the rigid carbon skeleton of sterols is particularly suitable to generate a very large number of derivatives, which differ by the carbon number and/or the position of various substituents (mainly hydroxyl or keto groups) (1). In addition to hormones, sterols give rise to bile acids/alcohols, initially considered as emulsifiers facilitating lipid digestion, but nowadays also known as important signalling molecules acting on specific receptors (2).

### Diversity of steroid mechanisms of action

Our concepts on (steroid) hormone mechanism of action has evolved. In the classical scheme, steroid hormones interact with nuclear receptors and the complex formed regulates the transcriptional activity of target genes, which promoters contain specific sequences (hormone-responsive elements). But steroids also act at cell membrane level where they elicit rapid non-transcriptional effects. Among the identified steroid membrane receptors, we may mention vertebrate GPER1/GPR30 (a membrane estrogen receptor – (3), TGR5 (a bile acid membrane receptor – (4), MARRS (a calcitriol receptor –(5) or drosophila DopEcR (a dopamine and ecdysone membrane receptor – (6). All these receptors belong to the family of GPCR/7TD receptors. Moreover, intact or truncated forms or the steroid nuclear receptors are bound to the plasma membrane and do not act there as transcription factors(7,8).

### Ecdysteroid effects on vertebrates

We are especially interested by the pharmacological effects of ecdysteroids on mammals. Ecdysteroids are a large family of steroids initially discovered in arthropods (zooecdysteroids) and later in plants (phytoecdysteroids) (9). They are present in many plant species where they can reach concentrations of up to 2-3 % of the plant dry weight, and they are expected to protect plants against phytophagous insects.

With the aim to use these molecules for crop protection, toxicological studies were performed on mammals, which unexpectedly concluded to both their lack of toxicity (oral LD_50_ > 9 g/kg) and their “beneficial” effects, *e.g*. anti-diabetic and anabolic properties (10). Such effects have to be linked with the presence of large amounts of phytoecdysteroids in several plants used worldwide by traditional medicine. At the moment, numerous effects have been reported, allowing to consider ecdysteroids as some kind of “universal remedy”(11). While many effects have been described on animals (10,12,13), the clinical evidence for 20E effectiveness in humans remains limited at the moment (14-16).

### How do ecdysteroids work?

In spite of more than 40 years of research, the mechanism of action of these molecules on mammals/humans has not been elucidated, as only diverging reports are available for the moment. Several data favour an action on membranes through a GPCR receptor(17), whereas other ones suggest the involvement of a nuclear receptor, the estrogen receptor ERβ (18,19).

There is in fact no direct evidence for the binding of 20E to nuclear estrogen (or androgen) receptors (12,13). The evidence of ERβ involvement in 20E effect is based on the use of specific pharmacological activators or inhibitors of ERs, the former being able to mimic and the latter to inhibit the effects of 20E on target cells such as osteoblasts (20) or myoblasts (19). These studies however do not bring proofs for a direct 20E binding, as ER receptor could be activated indirectly, and even in the absence of ligand, *e.g*. by phosphorylation (21). Some evidence for a direct binding was provided by *in silico* modelling (22), but this certainely does not represent a definite proof, as the result may strongly depend on the model used, and an opposite conclusion was drawn by Lapenna *et al*. (23).

Evidence for membrane effects of 20E is based on early studies showing the rapid modulation of several second messengers (cAMP, cGMP, IP3, DAG, Ca^2+^) in target cells (24-26) and on the fact that 20E bound to metallic nanoparticles, preventing its entrance in target cells, is still active (27). More recently, Gorelick-Feldman *et al*. (17) used a pharmacological approach with various inhibitors (*e.g*. pertussis toxin). They concluded that the membrane 20E receptor belongs to the GPCR family and proposed a mechanism of transduction involving an unidentified GPCR and a membrane calcium channel (Supporting Fig. S1).

The above pharmacological arguments appear strong enough to consider that the cell membrane is (maybe not exclusively) a site of action of 20E. The present experiments have been undertaken in an attempt to identify the/one GPCR involved in 20E effects, and to understand its estrogen-like effects using gene silencing or different pharmacological approaches.

## MATERIALS AND METHODS

### Chemicals

Except otherwise mentioned, all the reagents and chemicals were from Sigma (Saint-Quentin Fallavier, France). Peptides such as angiotensin-(1-7), A779 (Asp-Arg-Val-Tyr-Ile-His-D-Ala) and A1 (Asp-Arg-Val-Tyr-Ile-His-D-Pro) were custom-prepared by the IBPS peptide synthesis platform (Sorbonne University, Paris, France). 20-Hydroxyecdysone (20E) was obtained from Chemieliva Pharmaceutical (Chongqing, China) or from Patheon (Regensburg, Germany) and had a purity of 96.5-97%.

### Preparation of HSA-conjugated 20-hydroxyecdysone

22-succinyl-20E was prepared according to Dinan *et al*.(28). Coupling the 20E derivative to human serum albumin (HSA) was performed according to a method provided by Dr J-P Delbecque (JP Delbecque and M. de Reggi, personal communication). The 20E-HSA conjugate (**Supporting Fig. S2**) was analyzed by mass spectrometry to determine the number of 20E molecules coupled to each HSA molecule. Analyses were performed in a 4700 MALDI TOF/TOF proteomics analyzer (Applied Biosystems). The 20E conjugate was studied in linear mode, positive ion mode. Laser acceleration is set at 20kV and the default laser fluency was set at 2,000 and modified according to the signal-to-noise quality. The matrix used was α-cyano-4-hydrocinnamic Acid (HCCA). Samples were prepared following the dried-droplet method, 1 µL of a mixture of 1 µL of matrix (10 mg/mL) and 1 µL of sample was spotted and dried with gaseous nitrogen. The mass shift of HSA around 5 kDa after coupling indicates that 9 molecules of 20E derivatives are coupled to each albumin molecule.

### Cell culture

The C2C12 mouse myoblast cell line (29) was purchased from ATCC (CRL-1772). Except otherwise mentioned, culture media, serum, antibiotics and supplements were from Life technologies (Villebon-sur-Yvette, France). All cultures contained 100 U/mL of penicillin, and 100 μg/mL of streptomycin and are maintained in a 5% CO_2_, 95% air humidified atmosphere at 37°C. For C2C12 proliferation, cells were maintained in DMEM medium containing 4.5 g/L glucose supplemented with 10% FBS. C2C12 cells were maintained at low passage (3-20 passages) for all experiments to maintain the differentiation potential of the cultures. Cell confluency was always kept below or equal to ∼80%. For all experiments, cells were first seeded at 30,000 cells per well in 24-well plates. To induce differentiation, C2C12 at ∼80% confluency in proliferation medium were shifted to DMEM medium supplemented with either 2% FBS or 2% horse serum.

### Protein synthesis (^3^H-leucine incorporation)

C2C12 cells were grown on 24-well plates at a density of 30,000 cells/well in 0.5 mL of growth medium (DMEM 4.5 g/L glucose supplemented with 10% fetal bovine serum). Twenty-four hours after plating, the differentiation induction into multinucleated myotubes was carried out in DMEM 4.5 g/L glucose containing 2% fetal bovine serum. After 5 days, cells were pre-incubated in Krebs medium 1 h at 37°C before being incubated in DMEM media without serum for 2.5 h in the presence of radiolabeled leucine (5 µCi/mL) and DMSO (control condition) or Insulin Growth Factor (IGF-1, 100 ng/mL) or 20E (0.1 – 0.5 – 1 – 5-10 µM). At the end of incubation, supernatants were discarded and cells were lysed in 0.1 N NaOH for 30 min. The cell soluble fraction-associated radioactivity was then counted using Wallac Microbeta 1450-021 TriLux Luminometer Liquid Scintillation Counter (Wallac EG&G, Gaithersburg, MD, USA) and protein quantification was performed using the colorimetric Lowry method.

### Myostatin and MAS gene expression assays

Cells were plated at a density of 30,000 cells per well in 24-well plates and were grown overnight in 5 % CO_2_ at 37°C. On day 5 of differentiation, treatments were carried out for 6 h. At the end of incubation, RNAs were extracted and purified using the RNAzol (Eurobio, Les Ulis, France). RNAs were converted into cDNAs with High-Capacity cDNA Reverse Transcription Kit (Applied Biosystems, ThermoFisher, Villebon-sur-Yvette, France) before performing a quantitative PCR using iTaq SybrGreen (Biorad, Marnes-la Coquette, France).

Q-RT-PCRs were then performed using a 7900HT Fast Real-Time PCR detection system (Applied Biosystems) and standard qPCR program (1 cycle 95°C 15 min followed by 40 cycles 95°C 15s and 60°C 1 min). QRT-PCR master mix contained the 100 ng cDNA samples and a set of primers at final concentration of 200 nM designed into two different exons and described below. The quality of RNA was checked using the nanodrop™ technology (ThermoFisher) when necessary.

The relative differences in gene expression levels between treatments were expressed as increases or decreases in cycle time [Ct] numbers compared to the control group where the [Ct] value of each gene was normalized to the beta actin gene or hypoxanthine guanine phosphoribosyl transferase (HPRT) gene. Results of gene expression were expressed in 2^-ΔΔCT^ after normalization with house-keeping genes. Primer sequences used are described in **Table 1**.

**Table 1.**
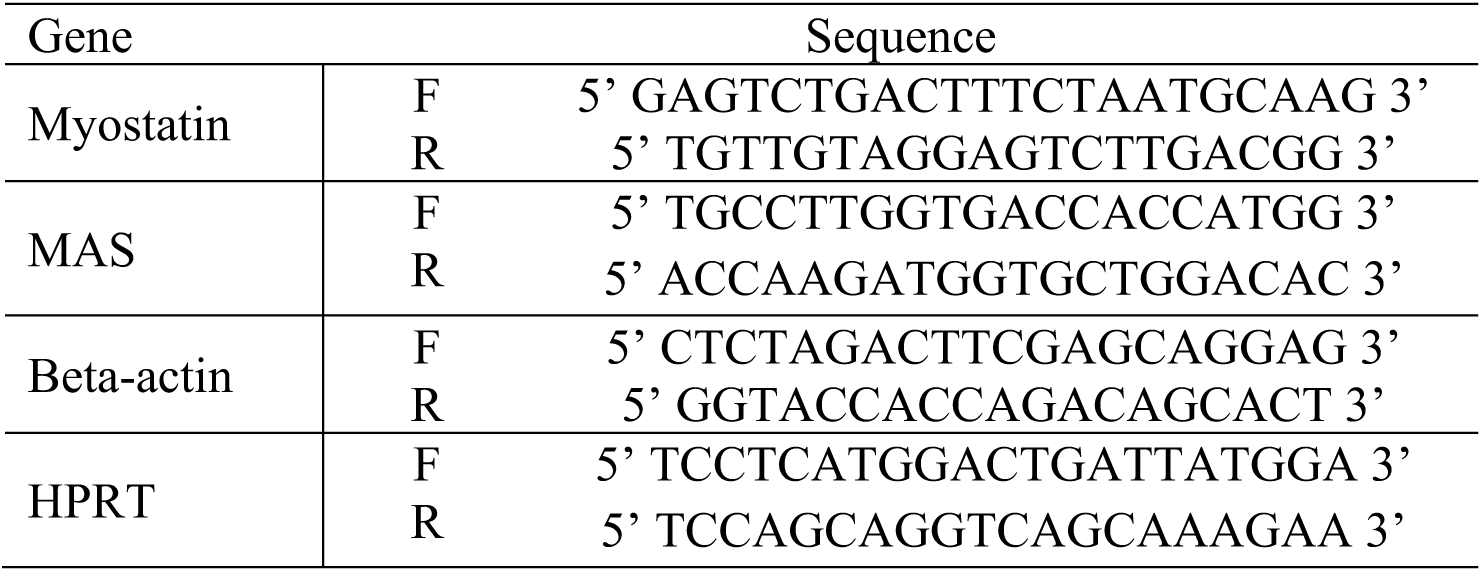
Primers used for mRNA quantification by RT-QPCR.

### SiRNA MAS assay

Cells were plated at a density of 10,000 cells per well in 24-well plates. After 3 days of differentiation, cells were transfected either with scramble SiRNA (10 nM) or MAS-1 SiRNA (10 nM) according to manufacturer’s instructions (Origene Technologies, Rockville, MD, USA). Two days after transfection, myotubes were treated with either DMSO or IGF-1 or 20E or angiotensin-(1-7) for 6 h. At the end of incubation, RNA was extracted and analyzed by QRT-PCR as described above.

### Binding studies

The affinity of 20E for human nuclear steroid receptors such as androgen receptor (AR), estrogen receptors alpha and beta (ERα, ERβ) and glucocorticoid receptor (GR) was determined by radioligand binding assays (CEREP/Eurofins). The selective ligands [^3^H]-methyltrienolone, [^3^H]-estradiol, [^3^H]-dexamethasone were employed on cells expressing either human endogenous or recombinant AR, ERα or β, or GR, respectively. 20E was used at concentrations up to 100 μM as a potential competitor. Inhibition of control specific binding was determined, and IC_50_ and K_i_ were calculated when possible. Additionally, a receptor screen was carried out on 45 GPCR and 5 nuclear receptors at a fixed concentration of 20E (10 μM). Radioligand binding assays were performed according to manufacturer instructions employing ^3^H- or ^125^I-labelled specific ligands of each receptor (SafetyScreen87 Panel, Panlabs, Taipei, Taiwan).

### Statistical analyses

Statistical analysis was performed using Graph Pad Prism® Software. Anova followed by a Dunnett t-test or a Kruskal Wallis followed by a Dunn’s test when the variances significantly differed have been performed. To evaluate the significance of differences between two groups, the choice of parametric Student t-test or non-parametric Mann-Whitney test was based on the normality or non-normality of data distribution, respectively (D’Agostino & Pearson test). The results are considered significant at p-value <0.05 (*), <0.01(**), <0.001 (***).

## RESULTS

### 20-Hydroxyecdysone stimulates muscle anabolism

20-Hydroxyecdysone (20E) effects were investigated on pre-established murine myotubes (following 6 days of differentiation). An anabolic effect was investigated through *de novo* protein synthesis assay. A dose-dependent increase in protein synthesis was observed in response to 20E treatment of C2C12 myotubes versus untreated conditions (**Fig. 1A**). IGF-1 (100 ng/mL), employed as positive control (30) displayed, as expected, an improvement in [^3^H]-Leu incorporation (+20%, p < 0.001). 20E effect was significant from 0.5 μM to 5 μM. The maximal effect (+ 27 %; p < 0.001) was measured with 5 μM of 20E, while a treatment with a higher concentration of 20E (10 μM) appeared to be notably less efficient (+11%, ns) than the previous concentration tested.

**Figure 1:**
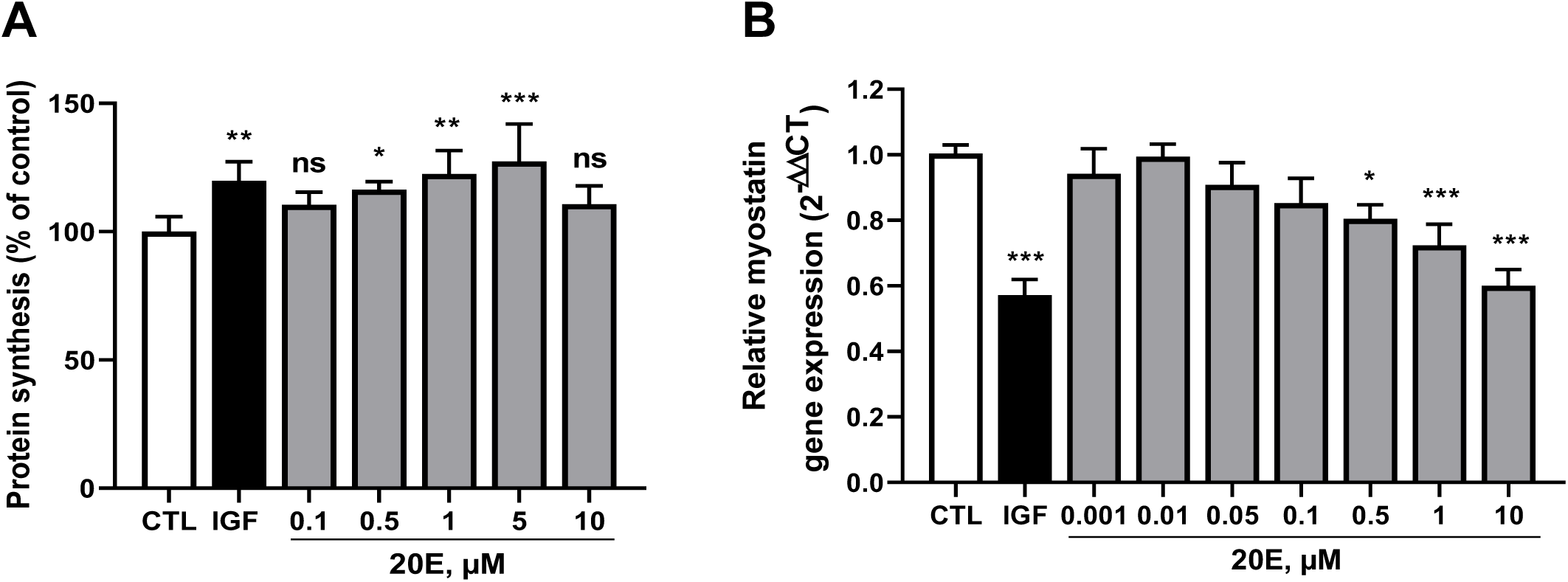
Effects of 20E on protein synthesis and Myostatin gene expression in C2C12 cells. **(A)** Experiments displaying 20E effects on protein synthesis in differentiated myotubes detected by [3H]-leucin incorporation. Results are shown as means ± SEM, ***p < 0.001, **p < 0.01, *p < 0.05 vs untreated control (Kruskal-Wallis followed by a Dunn’s test). (**B**) C2C12 mouse myoblasts were differentiated for 6 days into myotubes. They were then treated for 6 hours with concentrations of 20E ranging from 0.001 to 10 μM. Myostatin gene expression was detected by qRT-PCR. Results are shown as means ± SEM with ***p<0.001, **p<0.01, *p<0.05 vs untreated control (one-way ANOVA with Dunnett’s test compared to untreated control).

Myostatin is a major autocrine regulator that inhibits muscle growth in mammals. The myostatin transcript bioassay was developed and standardized in order to assess ecdysteroid activity (31). IGF-1 (100 ng/mL) used as a positive control demonstrated a significant inhibition of myostatin gene expression (57% of untreated control cells, p<0.001, **Fig. 1B**). A dose-dependent and partial inhibition of myostatin gene expression was observed in response to 20E treatment at concentrations comprised between 0.001 and 10 µM. This inhibition reached significance from 0.5 µM 20E (**Fig. 1B**). The inhibition of myostatin gene expression was then employed as a readout for 20E activity.

### 20-Hydroxyecdysone acts on cell membranes

In order to confirm that 20E acts primarily on cell membrane, or if it needs to penetrate into the cell to exert its effects, a membrane-impermeable derivative of 20E was produced (**Supporting Fig. S2**). We compared the effects of free 20E and its 22S-HSA conjugate on myostatin gene expression at the concentration of 10 µM 20E-equivalents (**Fig. 2**). We observed that the membrane-impermeable 20E-derivative retained an activity similar to that of free 20E. This results demonstrates that the presence of a bulky protein does not prevent 20E activity and is an additional argument for the interaction of 20E with a membrane receptor, as proposed by Gorelick-Feldman *et al*. (2010).

**Figure 2:**
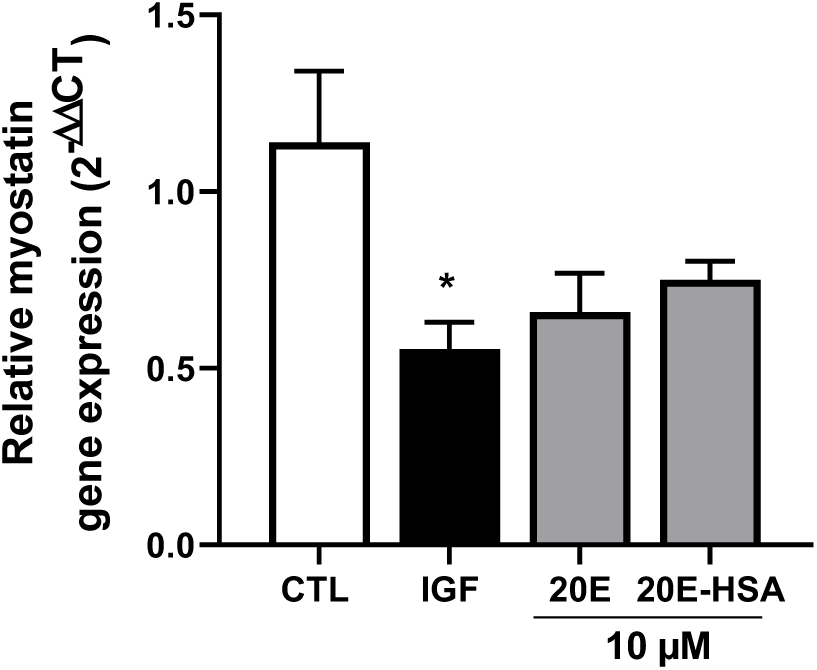
20E acts on C2C12 myotubes from outside of the cell. C2C12 mouse myoblasts were differentiated for 6 days into myotubes. They were then treated for 6 hours with IGF-1 (100 nM), 20E (10 μM) or 20E-HSA (10 μM 20E-equivalent). Myostatin gene expression was detected by qRT-PCR. Results are shown as means ± SEM with *p < 0.05 vs untreated control (one-way ANOVA with Dunnett’s test compared to untreated control).

### 20-Hydroxyecdysone acts via a GPCR-type receptor

The GPCR hypothesis is based on the inhibition of 20E effects by pertussis toxin, but there are still numerous possible GPCR candidates. We selected a set of GPCR receptors based on available literature and using different criteria corresponding to well-established effects of 20E: (i) involvement in the control of muscle cells activity and glycaemia/insulin sensitivity, and (ii) ability to reduce fat mass gain in high-fat fed animals (32,33). A set of GPCRs was thus selected including TGR5 (bile acids receptor - (4), GPER/GPR30 (estradiol, aldosterone receptors -(3,6), LPA1 (lysophosphatidic acids receptor -(34), APJ (apelin receptor -(35), OXTR (oxytocin receptor -(36), AVPR1 (vasopressin receptor -(37), MrgD (alamandine receptor - (38), MARRS (vitamin D3 receptor - (5) and Mas (angiotensin-(1-7) receptor -(39). The possible interaction of 20E with those receptors was assayed using different approaches according to available methodologies: (1) *in silico* binding when 3D structures were available, (2) *in vitro* direct binding studies by competition with a radioactive natural ligand, (3) comparison of the effects of agonists with those of 20E on C2C12 cells or (4) effect of known antagonists on the response of C2C12 cells to 20E. Using this approach, the only receptor which gave positive data was Mas, the receptor of angiotensin-(1-7).

### 20-Hydroxyecdysone and Angiotensin-(1-7) act via Mas receptor activation

Using the myostatin gene expression assay, a pharmacological approach was emloyed to compare the effects of the endogenous Mas receptor agonist (angiotensin-(1-7)) with those of 20E on C2C12 cells in the presence and absence of known antagonists.

Angiotensin-(1-7) (Ang 1-7, **Supporting Fig. S3**) as well as 20E (**Fig. 1B**) partially inhibits myostatin gene expression in a dose-dependent manner. As expected, this inhibition by Ang 1-7 (10 µM) was totally abolished by specific Ang 1-7 antagonists (A1 or A779, 10 µM) (**Fig. 3A**). Interestingly, 20E inhibitory effects on myostatin gene expression was also fully reverted by the same antagonists (**Fig. 3A**) suggesting that inhibition of myostatin gene expression by 20E (or Ang 1-7) is mediated by the receptor of Ang 1-7. In contrast, the effect of IGF-1, which acts through its own receptor (insulin-like growth factor 1 receptor; IGF1R) remains unchanged in the presence or in the absence of Ang 1-7 antagonist (**Supporting Fig. S4**).

**Figure 3:**
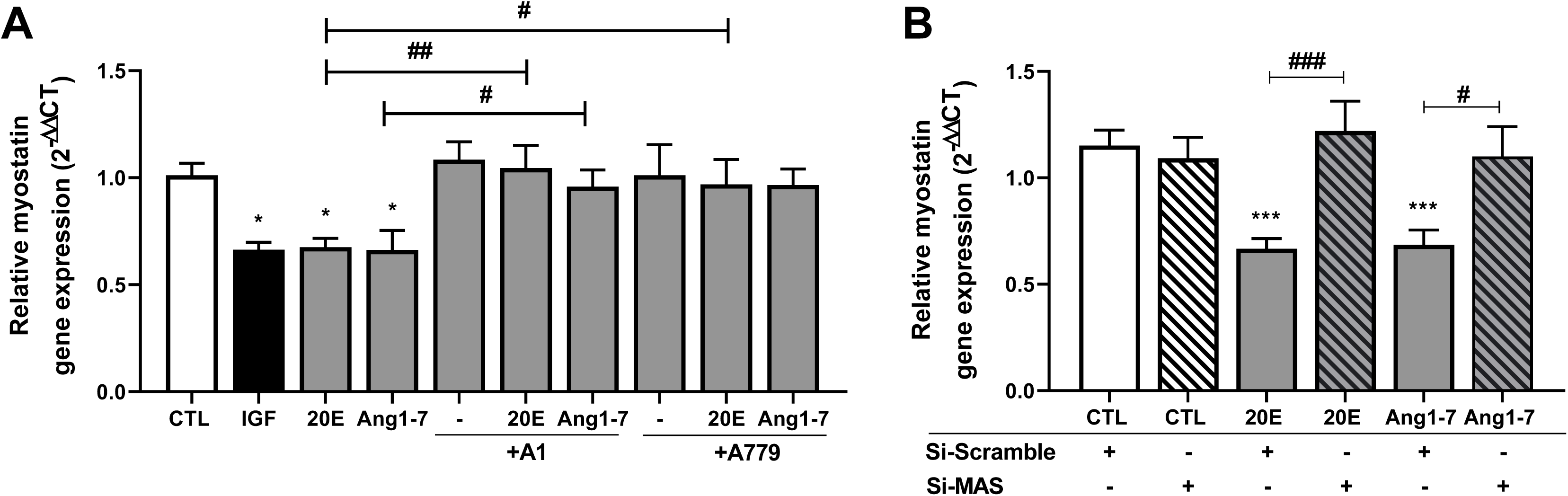
20-Hydroxyecdysone effects are mediated through Mas receptor activation. C2C12 mouse myoblasts were differentiated for 6 days into myotubes. (**A**) Mas antagonists (A1 and A779) blocked 20E and Ang 1-7 inhibitory activities on myostatin gene expression determined by qRT-PCR. **(B)** MAS siRNA abolished 20E- and Ang 1-7-induced myostatin inhibition. Results are shown as means ± SEM with *p < 0.05; **p < 0.01; ***p < 0.001. ANOVA with Dunnett’s test compared to untreated control and Mann-Whitney test was performed to compare two groups with # p<0.05; ## p<0.01, ### p<0.001.

In order to further assess the involvement of Mas receptor in 20E activity on myotubes, we designed a gene interference experiment using silencing RNA directed against Mas receptor (SiRNA MAS). The efficiency of Mas receptor down regulation by SiRNA was tested first. A significant decrease of MAS gene expression by a factor 2 in all transfected group by directed SiRNA versus scramble SiRNA was observed (**Supporting Fig. S5**). In a similar way to what was observed with antagonists (**Fig. 3A**), down regulation of Mas receptor reversed 20E or Ang1-7 effects on myostatin gene expression (**Fig. 3B**). As expected, and consistently with the pharmacological approach, down regulation of Mas receptor had no impact on the effect of IGF-1 on myostatin gene expression (data not shown).

### 20-Hydroxyecdysone does not bind to a set of nuclear receptors

Parr *et al*. (19,22) have proposed that 20E effects are explained through its binding to estrogen receptor ERβ. It is however difficult to consider that this receptor corresponds to a canonical nuclear receptor of estradiol, given that several binding studies to nuclear ERs were unsuccessful (12,13,17). We too performed binding tests of 20E for AR, ERα and ERβ which were all negative at up to 100 µM (data not shown). Similarly, an off-target safety screen for 87 receptors was equally negative for ERα and AR (**Supporting Table S1**). Nevertheless, Parr’ results are not unique. Thus Gao *et al*. (20) showed that 20E can activate several ERβ target genes, and indeed there are multiple similarities between the effects of 20E and 17β-Estradiol (E2) on muscles (40) or skin cells(41). Thus, while the above binding studies seem to exclude canonical nuclear forms of ERs, the question remains open for the membrane ones.

### Estradiol effects on C2C12 myotubes

In an attempt to explain this discrepancy, we engaged a set of experiments to characterize estradiol effects on C2C12 cells and to identify which type of E2 receptor could be involved in 20E effects. We first checked if, like 20E, E2 was able to impact myostatin gene expression in C2C12 cells. Myotubes were treated with increasing doses of E2 during six hours and myostatin expression was then determined at transcriptional level. E2 significantly inhibits myostatin gene expression from 0. 1 µM to 1 µM (**Fig. 4A**). To determine if E2 activity relies on an interaction with a plasma membrane receptor, we employed the same strategy as the one presented above for 20E (**Fig. 2**). We used a membrane-impermeable conjugate of E2 made of bovine serum albumin (BSA) and a carboxymethyloxime (CMO) linker. E2-CMO significantly inhibited myostatin gene expression (−39%, p < 0.05) with the same trend as IGF-1 positive control (−47%, p < 0.01). By contrast, E2-CMO-BSA-conjugate was inactive (**Fig. 4B**) and did not significantly decrease myostatin expression (−8 %, ns). This experiment allows to exclude an interaction of E2 with a transmembrane receptor, but rather could possibly fit with a receptor bound to the cytoplasmic side of the cell membrane by a lipid anchor.

**Figure 4:**
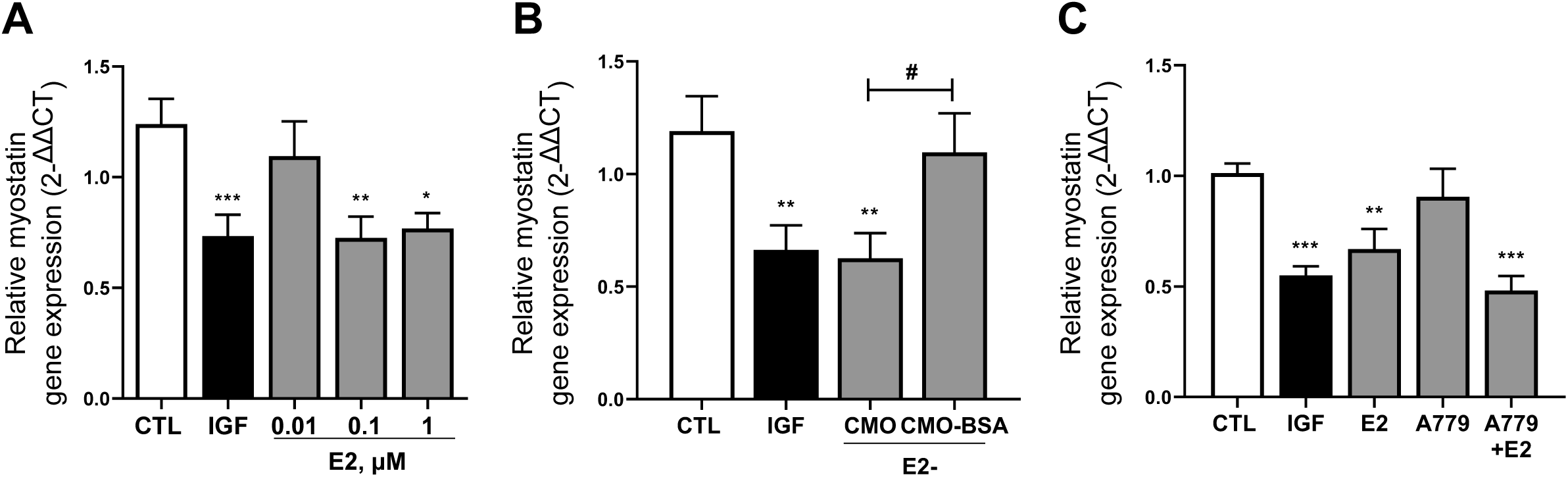
17β-Estradiol-mediated myostatin inhibition is not linked with a transmembrane receptor and is not blunted by a Mas antagonist. (**A**) Differentiated C2C12 cells were treated with IGF-1 (100 ng/mL) or 17β-estradiol (E2; 0.01, 0.1 and 1µM) for 6 h. Myostatin gene expression was analysed by qRT-PCR. (**B**) Effect of E2-CMO (0.1 µM) and E2-CMO-BSA (0.1 µM) on myostatin gene expression (qRT-PCR). (**C**) Effect of E2 (0.02 µM) in combination with Mas antagonist A779 (10 µM) on myostatin gene expression. Results are shown as means ± SEM with *p < 0.05; **p < 0.01; ***p < 0.001 compared to untreated control; house-keeping gene used was HPRT.

Sobrino and colleagues (42) showed that the vasodilatory effect of E2 was blunted by a Mas antagonist. This engaged us to check if a Mas antagonist would inhibit the effect of E2 on MSTN gene expression by C2C12 cells. E2 effect on myostatin was tested in the presence of a Mas antagonist (A779) but, unexpectedly, the effect of E2 was not inhibited by A779 (**Fig. 4C**).

*17α-E2* is an epimer of estradiol that does not bind nuclear ERα or ERβ and is known to bind only membrane forms of ER (43). This compound proved active for the inhibition of MSTN gene activity (**Supporting Fig. S6**). This result provides an additional argument for the involvement of a non-nuclear ER.

## DISCUSSION

Our data combined with those previously available from the literature allow us to conclude that the effects of 20E on C2C12 cells involve both Mas receptor and a non-nuclear estradiol receptor. Although these results do not allow to identify unambiguously its primary target, they allow to reconcile the findings of Gorelick-Feldman *et al*. (17) and Parr *et al*. (19) thanks to a mixed mechanism of action.

Indeed, of paramount importance is our finding that a membrane impermeable form of E2 is inactive, whereas a membrane-impermeable form of 20E still remains active. This allows us to conclude that 20E does not bind the concerned ER receptor, and that the activation of this estrogen receptor by 20E is secondary to Mas activation.

Five different E2 receptors have been described (not including splice variants) including 2 GPCRs(44): GPR30(45), and Gq-mER (46) – plus possibly another plasma-membrane associated receptor, ER-X(47). Our experiments with GPR30 agonists and antagonists exclude GPR30 from the candidates. In addition, nuclear canonical receptors being excluded, the question remains open regarding which membrane E2 receptor form is involved in the effects of E2 and 20E on muscle cells.

Different membrane forms of the nuclear estrogen receptors (ERα and ERβ) produced by alternative splicing have been described, that may correspond to either palmitoylated full forms or truncated forms unmasking a potential transmembrane alpha-helix sequence(7,8,48). Non-truncated ERs may reversibly bind to the cytoplasmic face of the membrane by a S-palmitoyl anchor (49).

In this respect, experiments on C2C12 cells treated with diarylheptanoid compounds (HPPH) provide very important informations(50,51). These molecules display growth- and differentiation-promoting effects on C2C12 cells that involve a membrane form of ERα receptor bound by a palmitoyl anchor, and they were abolished by 2-bromohexadecanoic acid, an inhibitor of palmitoylation(50).

Interestingly, Garratt *et al*. (52) have shown that 17αE2, an E2 epimer that does not bind nuclear forms of estrogen receptors has beneficial effect on muscles, notably during sarcopenia. These results demonstrate that nuclear and membrane forms of E2 receptors display different ligand specificities. Therefore, the use of “selective” inhibitors based on their effects on nuclear ERs does not allow unambiguous conclusions about whether Erα or ERβ membrane forms are targeted, and specific silencing experiments have to be preferred.

Membrane forms of ERs can associate with a GPCR: for example, in neuronal membranes, ER is associated with mGLUR1a (a glutamate receptor) and in this system, E2 effects are blunted by a mGLUR1a antagonist(53).

A functional interaction between ER and Mas receptor has already been observed by Sobrino *et al*. (42) in HUVEC cells. These authors observed that E2 effect (increased NO synthesis) was abolished by an antagonist of Mas receptor. An association between Mas and a non-nuclear form of ER seems therefore highly probable.

Taken together, our data best fit with the presence of a complex associating Mas and a estrogen membrane receptor (**Fig. 5**), likely together with a caveolin and/or striatin, as well as additional transduction effectors (e.g. Gq, NOS, …). The demonstration of such a functional association will require membrane fractionation techniques combined with various immunoprecipitation techniques. Whether a symmetrical interaction of this ER and IGFR exists is an attractive possibility, which is presently under investigation.

**Figure 5:**
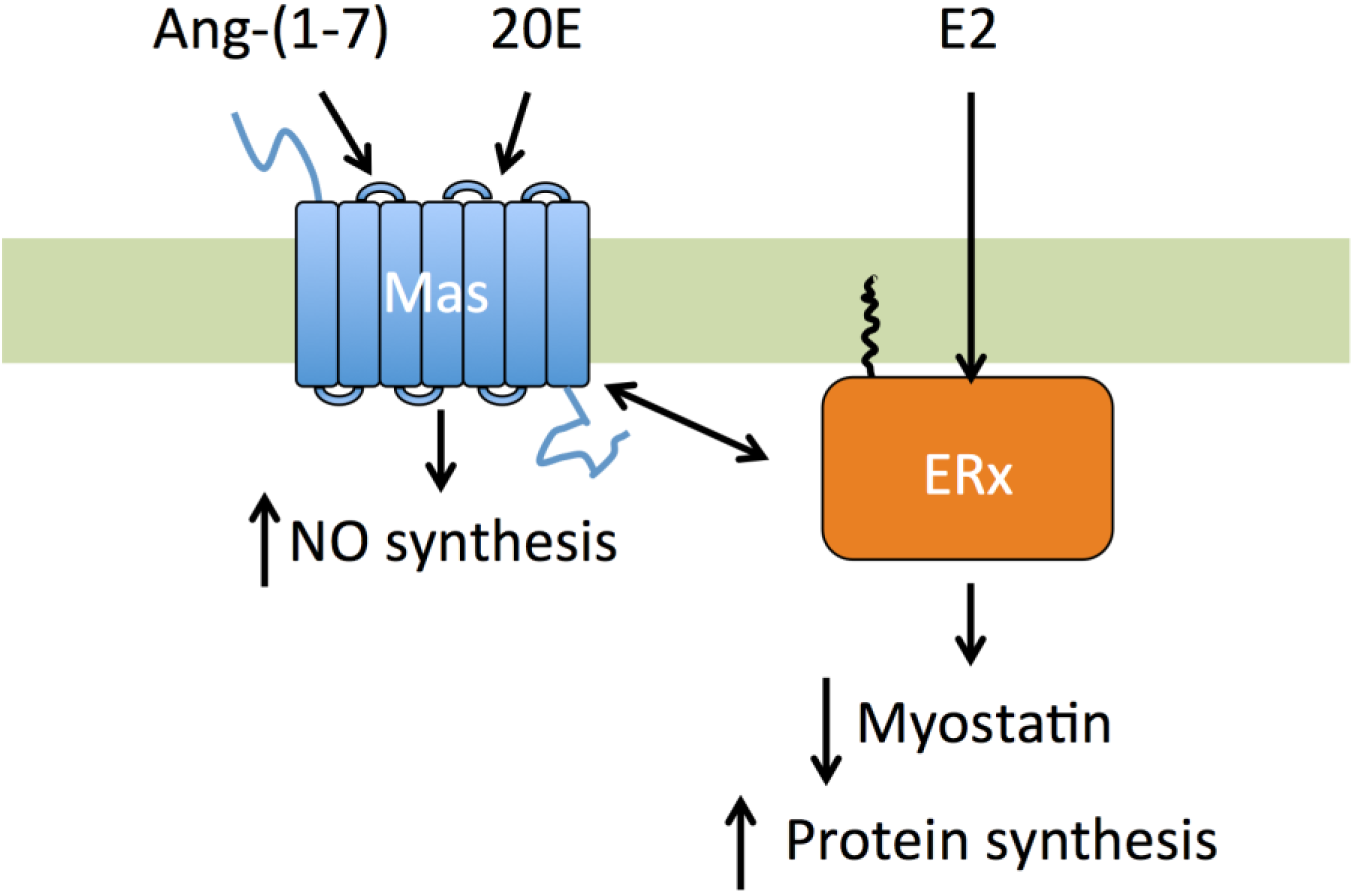
Proposed mechanism of MSTN gene control by Ang-(1-7), 20E and E2. According with this scheme, myostatin and protein synthesis would be controlled directly by ER and indirectly by Mas, while NO synthesis would be controlled directly by Ang-(1-7) (Tirupula *et al*., 2015) or 20E (Omanakuttan *et al*., 2016), and indirectly by E2 (Sobrino *et al*., 2017).

The case of ER is not unique. Ruhs *et al*., (54) described an association between aldosterone receptor (MR) and GPER/GPR30: accordingly, some effects of aldosterone are inhibited by G15, a GPER inhibitor (55) and, most interestingly, they also showed an association between MR and the angiotensin II receptor AT1. Interestingly, we would thus be in the presence of both a complex between aldosterone and angiotensin II AT1 receptors and, symmetrically, of a complex between estradiol and angiotensin-(1-7) Mas receptors and displaying opposite physiological effects.

It is worth mentioning that 20E will only activate a particular membrane form of ER, whereas E2 would in addition bind nuclear receptor(s). Thus 20E is devoid of any feminizing activity (13,56) and is probably inactive on estrogen-dependent cancer cells.

## CONCLUSION

It is noteworthy that both Ang-(1-7), E2 and 20E display similar pleiotropic effects on many different organs/functions (**Supporting Table 2**), During the recent years, the number of beneficial effects of the protective arm of the renin-angiotensin system has been continuously increasing, and from the available data, it is expected that 20E will provide similar beneficial effects on several types of diseases (sarcopenia, diabetes, metabolic syndrome etc.).

Based on these above findings, a pharmaceutical grade preparation of 20E (BIO101) has been developed and is presently being assayed in a phase 2 clinical trial for treating sarcopenia, (a double-blind, placebo controlled, randomized interventional clinical trial (SARA-INT), ClinicalTrials #NCT03452488). We are confident that the beneficial effects of 20E/BIO101 will also be further established on e.g. lungs, kidneys and cardiovascular pathologies and could offer new therapeutic strategies.

## Acknowledgements

Dr JP Delbecque (University of Bordeaux) for his help for the synthesis of 20E-HSA conjugate and Dr L. Dinan for language improvement and critical reading of the manuscript

## Abbreviations

(20E): 20-Hydroxyecdysone,
(17α-E2): 17α-Estradiol,
(E2): 17β-Estradiol,
(HCCA): α-cyano-4-hydrocinnamic acid,
(AR): androgen receptor,
(Ang1-7): angiotensin-(1-7),
(AT1R): angiotensin II receptor type 1,
(A779): Asp-Arg-Val-Tyr-Ile-His-D-Ala,
(A1): Asp-Arg-Val-Tyr-Ile-His-D-Pro,
(BSA): bovine serum albumin,
(CMO): carboxymethyloxime,
(cAMP): cyclic adenosine monophosphate,
(cGMP): cyclic guanosine monophosphate,
(DAG): diacylglycerol,
*(DMSO)*: *dimethyl sulfoxide*,
(*DopEcR*): dopamine/ecdysteroid *receptor*,
*(DMEM)*: *Dulbecco’s Modified Eagle Medium*,
(EcR): ecdysone receptor,
(ERβ): estrogen receptor β,
(ERα): estrogen receptor α,
(ERs): estrogen receptors,
(GR): glucocorticoid receptor,
(GPCR): G protein-coupled receptor,
(GPR30): G protein-coupled receptor 30,
(GPER): G protein-coupled estrogen receptor 1,
(*TGR5)*: G-protein-coupled bile acid receptor,
(Gq): G-protein subtype q,
(HSA): human serum albumin,
(HUVEC): human umbilical vein endothelial cells,
(HPRT): hypoxanthine guanine phosphoribosyl transferase,
(IP3): inositol trisphosphate,
(IGF-1): insulin growth factor,
(IGFR): insulin growth factor receptor,
(MALDI TOF/TOF): matrix-assisted laser desorption ionization - tandem time-of-flight,
(LD50): membrane-associated, lethal dose 50%,
(NO): nitric oxide,
(NOS): nitrous oxide synthase,
(QRT-PCR): quantitative reverse transcriptase polymerase chain reaction,
(*M*ARRS): rapid response steroid-binding *receptor*,
(7TD): seven transmembrane domains,
(siRNA): small interfering RNA,
([^3^H]-Leu): tritiated leucine,

---

**Supporting Figure S1:**
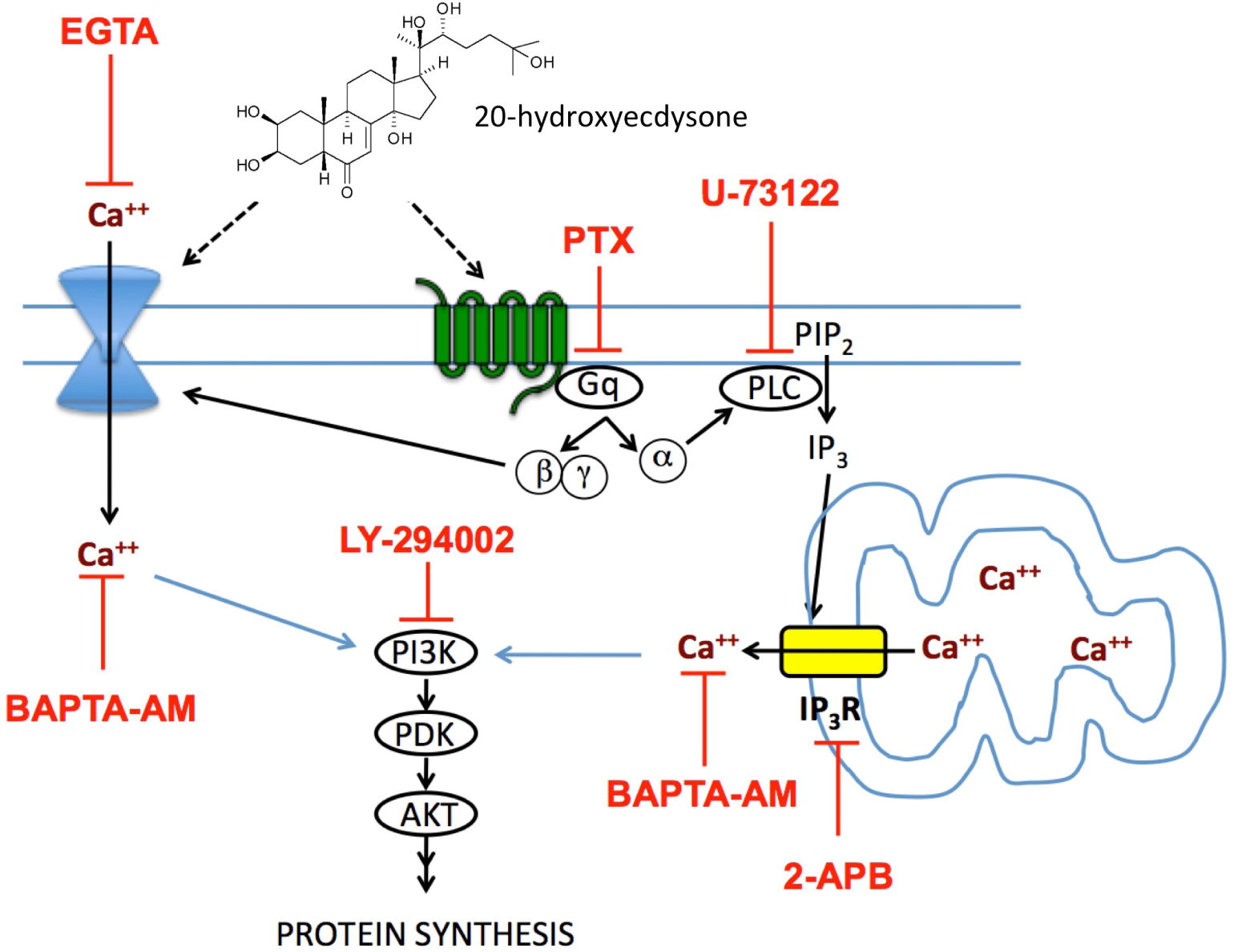
Proposed mechanism of 20E membrane action. (redrawn and modified from Gorelick-Feldman *et al*., 2010) Gq: a subtype of G protein; PLC: Phospholipase C; PIP_2_: Phosphatidylinositol 4,5-bisphosphate; IP_3_:Inositol triphosphate; IP_3_R: Inositol triphosphate receptor; PI3K: phosphoinositide 3-kinase, PDK: Pyruvate dehydrogenase kinase; AKT: Protein kinase B; EGTA: ethylene glycol tetraacetic acid, a calcium chelator; PTX: an inhibitor of G-protein coupled receptors; U-73122: an inhibitor of agonist-induced PLC activation; LY-294002: a potent inhibitor of phosphoinositide 3-kinases; BAPTA-AM: 1,2-bis(o-phenoxy)ethane-N,N,N’,N’-tetraacetic acid, a membrane permeable calcium chelator; 2APB: 2-Aminoethoxydiphenylborate, an inhibitor of IP_3_R and Transient Receptor Potential channel.

**Supporting Figure S2:**
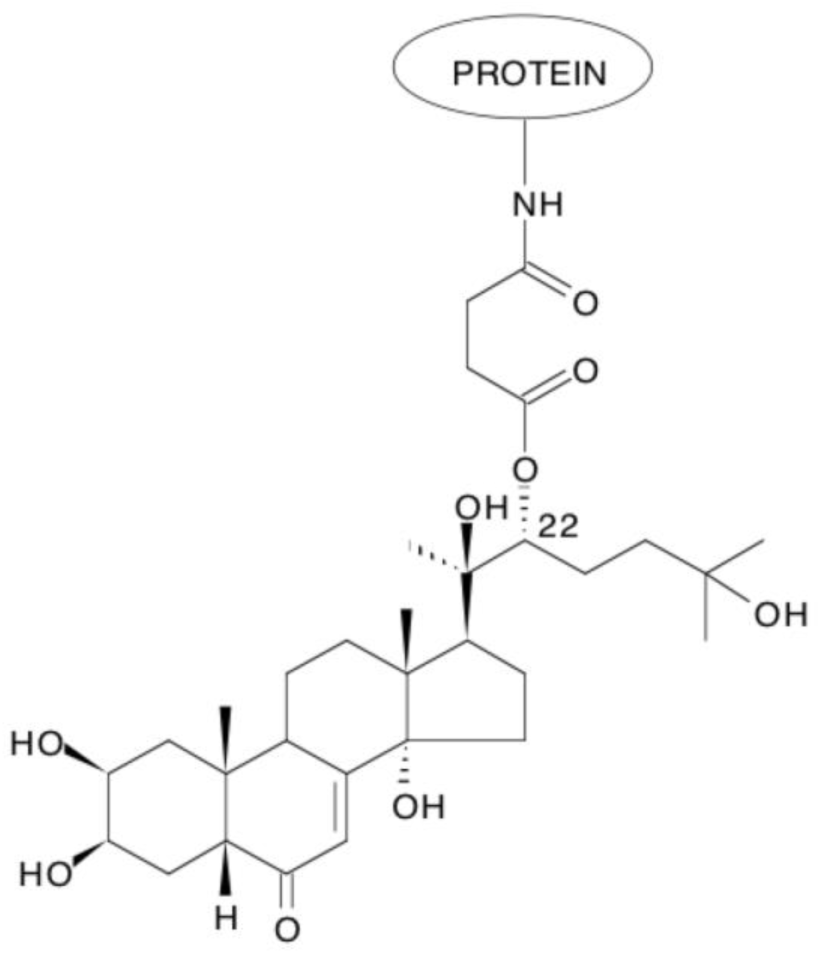
Structure of 20E 22-succinate coupled with human serum albumin.

**Supporting Figure S3:**
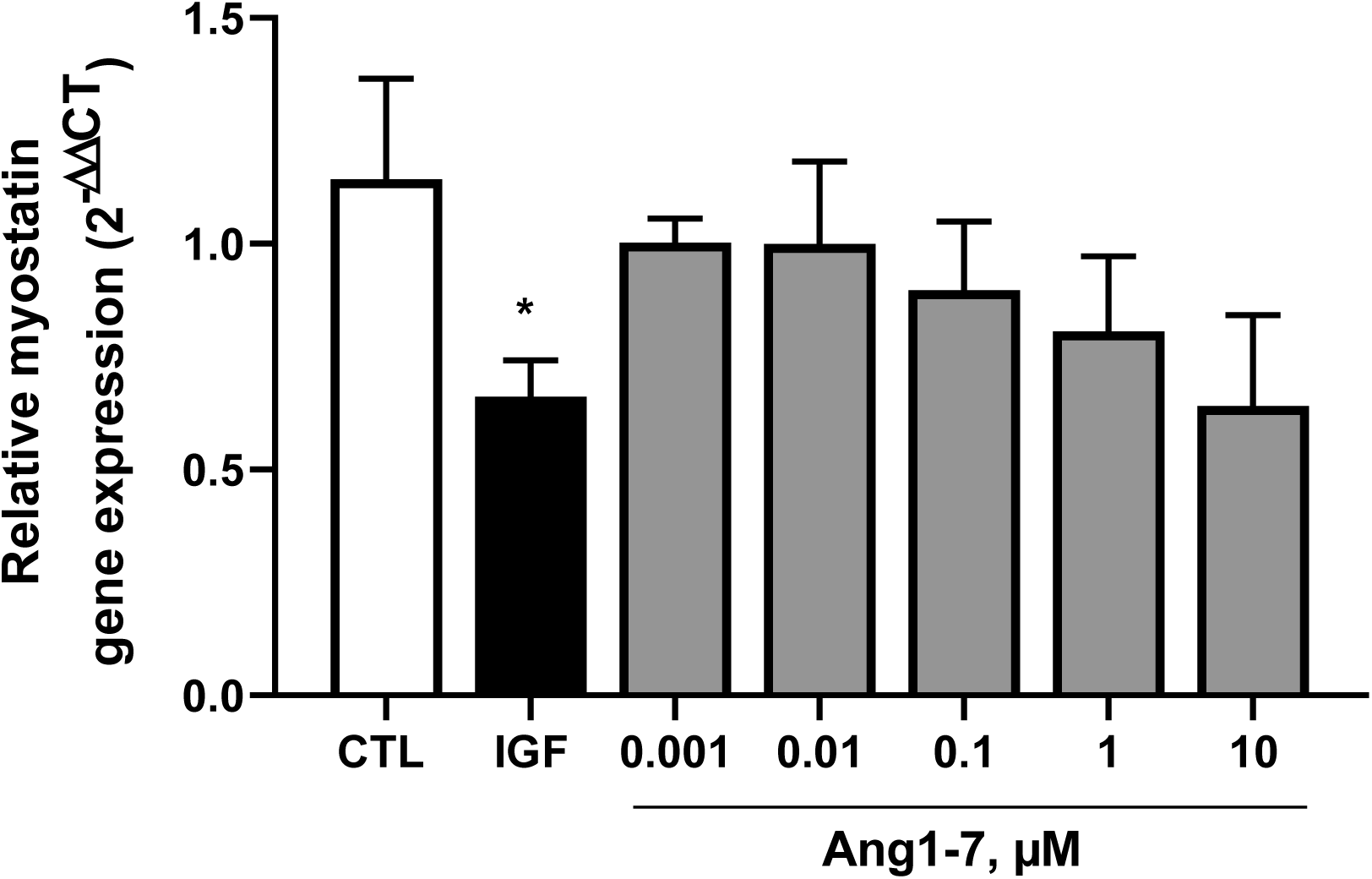
Effect of Angiotensin-(1-7) on myostatin gene expression. C2C12 mouse myoblasts were differentiated for 6 days into myotubes. They were then treated for 6 hours with concentrations of angiotensin 1-7 ranging from 0.001 to 10 μM. Myostatin gene expression was determined using qRT-PCR. Results are shown as means ± SEM with *p<0.05 (ANOVA with Dunnett’s test compared to untreated control).

**Supporting Figure S4:**
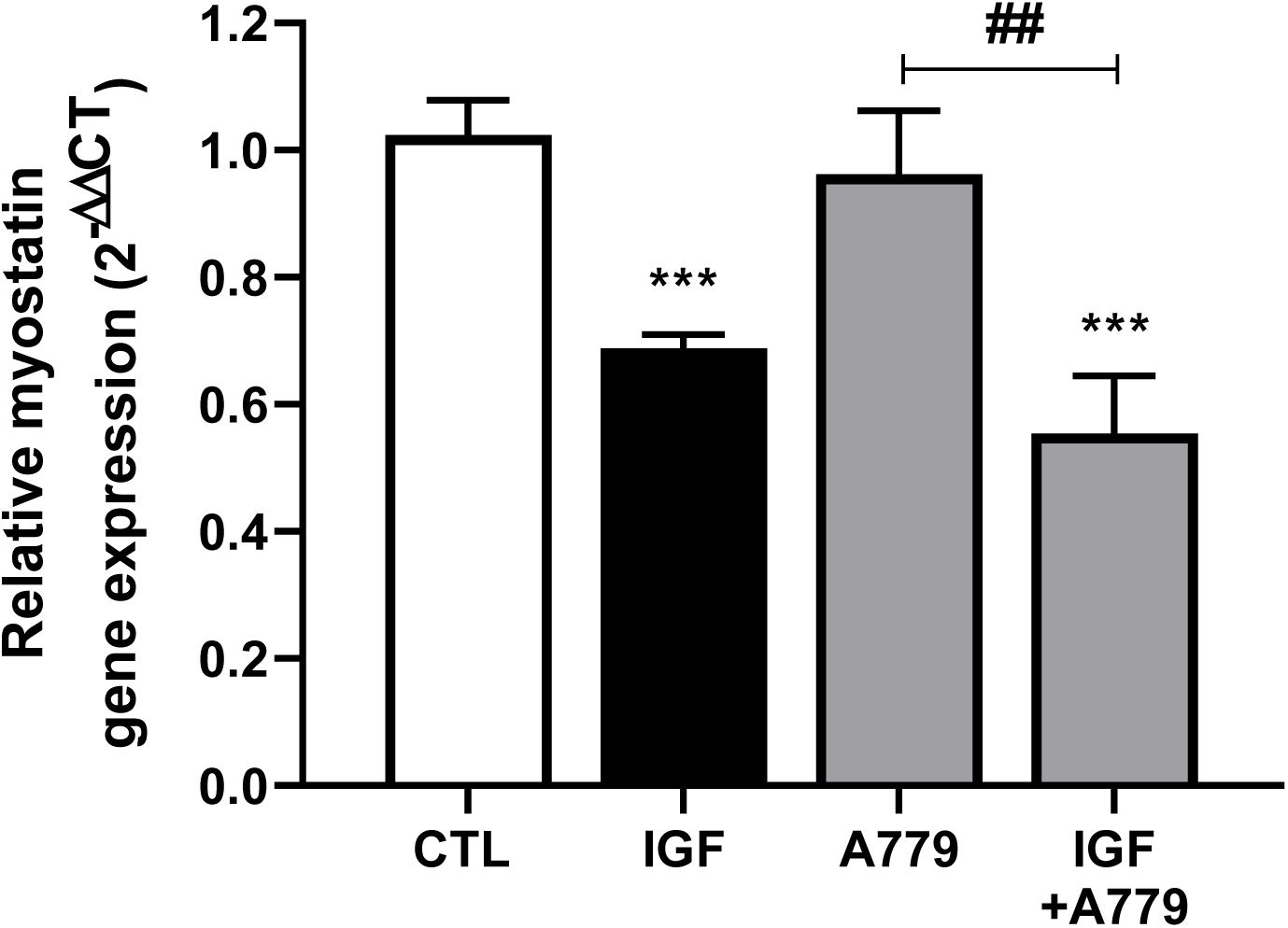
Effects of IGF-1 combined with A779 on myostatin mRNA expression. C2C12 mouse myoblasts were differentiated for 6 days into myotubes. IGF-1 (100 ng/mL) or vehicle was incubated for 6h with or without A779 (10 μM). Effects on myostatin gene expression were determined by qRT-PCR. Results are shown as means ± SEM with *p < 0.05, **p < 0.01; ***p < 0.001. D’Agostino and Pearson K2 test was employed to evaluated the normality followed by a Kruskal-Wallis test compared to untreated control. Mann-Whitney test was performed to compare two groups with ##p <0.01.

**Supporting Figure S5:**
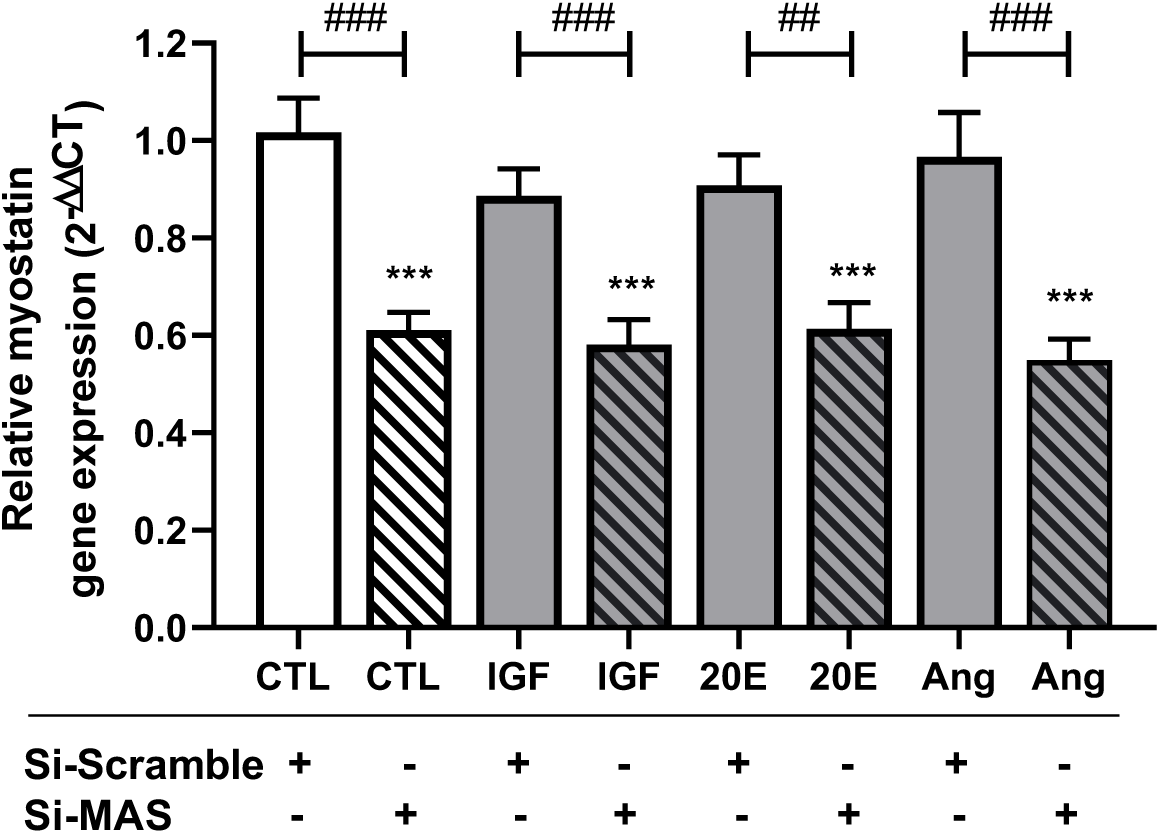
Inhibition of MAS expression by specific SiRNA. MAS mRNA expression in C2C12 transfected with scrambled siRNA (Si Scramble) or MAS-inhibiting siRNA (Si MAS) after 6 hours of exposure to 100ng/ml IGF-1, 10 µM 20E or 10 µM Angiotensin 1-7 (Ang). Results are shown as means ± SEM with ***p<0.001 ANOVA with Dunnett’s test compared to untreated control. Mann-Whitney test was performed to compare two groups with ## p<0.01, ### p<0.001.

**Supporting Figure S6:**
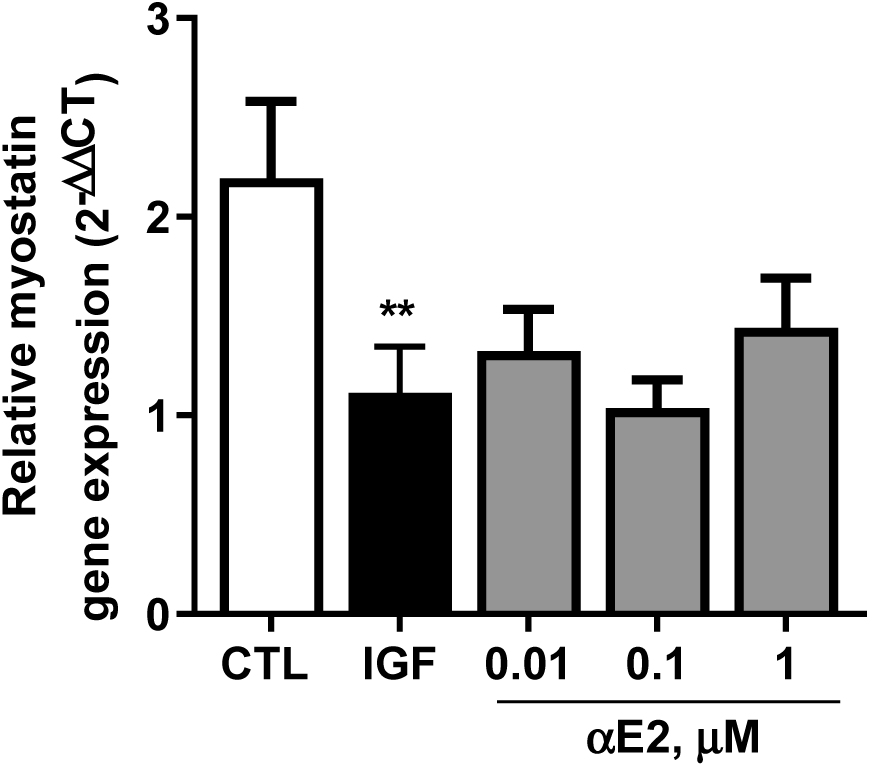
Inhibitory activity of 17α-estradiol (αE2) on myostatin gene expression. Differentiated C2C12 cells were treated with IGF-1 (100 ng/mL) or α-estradiol (0.01, 0.1 and 1 µM) for 6 h. Myostatin gene expression was analysed by qRT-PCR. Results are shown as means± SEM. House-keeping gene used was HPRT. Mann-Whitney test was performed to compare two groups with **p < 0.01 compared to untreated control.

**Supporting Table 1:** Effect of 20-hydroxyecdysone on 45 selected GPCR and 5 nuclear receptors. 20-Hydroxyecdysone was employed at a fixed concentration of 10 μM in radioligand binding assays involving specific ligands of a selection of GPCR and nuclear receptors. The percentage of specific ligand binding inhibition produced by 20E at 10 μM in duplicate experiments is presented. None of these inhibitions were considered significant.

| Receptor type | Receptor name | Species | Specific ligand | Inhibition (%) |
| --- | --- | --- | --- | --- |
| GPCR | Adenosine A<sub>1</sub> | hum | [<sup>3</sup>H] DPCPX | 9 |
| GPCR | Adenosine A<sub>2A</sub> | hum | [<sup>3</sup>H] CGS-21680 | -6 |
| GPCR | Adrenergic α<sub>1A</sub> | rat | [<sup>3</sup>H] Prazosin | 8 |
| GPCR | Adrenergic α<sub>1B</sub> | rat | [<sup>3</sup>H] Prazosin | -3 |
| GPCR | Adrenergic α<sub>1D</sub> | hum | [<sup>3</sup>H] Prazosin | 7 |
| GPCR | Adrenergic α<sub>2A</sub> | hum | [<sup>3</sup>H] Rauwolscine | -2 |
| GPCR | Adrenergic α<sub>2B</sub> | hum | [<sup>3</sup>H] Rauwolscine | 7 |
| GPCR | Adrenergic β<sub>1</sub> | hum | [<sup>125</sup>I] Cyanopindolol | -1 |
| GPCR | Adrenergic β<sub>2</sub> | hum | [<sup>3</sup>H] CGP-12177 | 4 |
| GPCR | Angiotensin AT<sub>1</sub> | hum | [<sup>125</sup>I] (Sar1, Ile8)-Angiotensin II | -4 |
| GPCR | Bradykinin B<sub>2</sub> | hum | [<sup>3</sup>H] Bradykinin | 0 |
| GPCR | Cannabinoid CB<sub>1</sub> | hum | [<sup>3</sup>H] SR141716A | 17 |
| GPCR | Cannabinoid CB<sub>2</sub> | hum | [<sup>3</sup>H] WIN-55,212-2 | 4 |
| GPCR | Chemokine CCR1 | hum | [<sup>125</sup>I] MIP-1α | -6 |
| GPCR | Chemokine CXCR2 (IL-8R<sub>B</sub>) | hum | [<sup>125</sup>I] IL-8 | -3 |
| GPCR | Cholecystokinin CCK<sub>1</sub> (CCK<sub>A</sub>) | hum | [<sup>125</sup>I] CCK-8 | 3 |
| GPCR | Cholecystokinin CCK<sub>2</sub> (CCK<sub>B</sub>) | hum | [<sup>125</sup>I] CCK-8 | -1 |
| GPCR | Dopamine D<sub>1</sub> | hum | [<sup>3</sup>H] SCH-23390 | 6 |
| GPCR | Dopamine D<sub>2L</sub> | hum | [<sup>3</sup>H] Spiperone | -12 |
| GPCR | Dopamine D<sub>2S</sub> | hum | [<sup>3</sup>H] Spiperone | -5 |
| GPCR | Endothelin ET<sub>A</sub> | hum | [<sup>125</sup>I] Endothelin-1 | 0 |
| GPCR | GABA<sub>B1A</sub> | hum | [<sup>3</sup>H] CGP-54626 | 0 |
| GPCR | Glutamate, Metabotropic, mGlu<sub>5</sub> | hum | [<sup>3</sup>H] Quisqualic acid | 27 |
| GPCR | Histamine H<sub>1</sub> | hum | [<sup>3</sup>H] Pyrilamine | 6 |
| GPCR | Histamine H<sub>2</sub> | hum | [<sup>125</sup>I] Aminopotentidine | -4 |
| GPCR | Leukotriene, Cysteinyl CysLT<sub>1</sub> | hum | [<sup>3</sup>H] LTD4 | 6 |
| GPCR | Melanocortin MC<sub>1</sub> | hum | [<sup>125</sup>I] NDP-α-MSH | -3 |
| GPCR | Melanocortin MC<sub>4</sub> | hum | [<sup>125</sup>I] NDP-α-MSH | -4 |
| GPCR | Muscarinic M<sub>1</sub> | hum | [<sup>3</sup>H] N-Methylscopolamine | 8 |
| GPCR | Muscarinic M<sub>2</sub> | hum | [<sup>3</sup>H] N-Methylscopolamine | -11 |
| GPCR | Muscarinic M<sub>3</sub> | hum | [<sup>3</sup>H] N-Methylscopolamine | -5 |
| GPCR | Muscarinic M<sub>4</sub> | hum | [<sup>3</sup>H] N-Methylscopolamine | 7 |
| GPCR | Neuropeptide Y Y<sub>1</sub> | hum | [<sup>125</sup>I] Peptide YY | -1 |
| GPCR | Nicotinic Acetylcholine | hum | [<sup>125</sup>I] Epibatidine | 6 |
| GPCR | Nicotinic Acetylcholine α1 | hum | [<sup>125</sup>I] α-Bungarotoxin | 8 |
| GPCR | Opiate δ<sub>1</sub> (OP1, DOP) | hum | [<sup>3</sup>H] Naltrindole | 3 |
| GPCR | Opiate κ (OP2, KOP) | hum | [<sup>3</sup>H] Diprenorphine | -2 |
| GPCR | Opiate μ (OP3, MOP) | hum | [<sup>3</sup>H] Diprenorphine | 10 |
| GPCR | Serotonin 5-HT<sub>1A</sub> | hum | [<sup>3</sup>H] 8-OH-DPAT | 5 |
| GPCR | Serotonin 5-HT<sub>1B</sub> | hum | [<sup>3</sup>H] GR125743 | -7 |
| GPCR | Serotonin 5-HT<sub>2A</sub> | hum | [<sup>3</sup>H] Ketanserin | 2 |
| GPCR | Serotonin 5-HT<sub>2B</sub> | hum | [<sup>3</sup>H] Lysergic acid diethylamide | 9 |
| GPCR | Serotonin 5-HT<sub>2C</sub> | hum | [<sup>3</sup>H] Mesulergine | 0 |
| GPCR | Tachykinin NK<sub>1</sub> | hum | [<sup>3</sup>H] Substance P | 7 |
| GPCR | Vasopressin V<sub>1A</sub> | hum | [<sup>125</sup>I] Phenylacetyl Tyr(Me)PheGlnAsnArgProArgTyr | 3 |
| Nuclear | Androgen (Testosterone) | hum | [<sup>3</sup>H] Methyltrienolone | 11 |
| Nuclear | Estrogen ERα | hum | [<sup>3</sup>H] Estradiol | 0 |
| Nuclear | Glucocorticoid | hum | [<sup>3</sup>H] Dexamethasone | 6 |
| Nuclear | PPAR<sub>γ</sub> | hum | [<sup>3</sup>H] Rosiglitazone | -4 |
| Nuclear | Progesterone PR-B | hum | [<sup>3</sup>H] Progesterone | 13 |

**Supporting Table 2:** The close similarity of Angiotensin (1-7), AVE 0991, 20E and estradiol effects.

| Effect | Angiotensin 1-7<br>(or ACE2 stimulation) | AVE 0991<br>(non-peptidic Mas agonist) | 20E and/or other ecdysteroids | Estradiol |
| --- | --- | --- | --- | --- |
| **Anabolic (muscle)** | Acuña *et al.*, 2014; Cabello-Verrugio *et al.*, 2015, 2017; Cisternas *et al.*, 2015; Abrigo *et al.*, 2016; Morales *et al.*, 2014, 2016 |  | Chermnykh *et al.*, 1988; Syrov, 2000; Tóth *et al.*, 2008; Gorelick-Feldman *et al.*, 2008, 2009, 2010; Lawrence, 2012; Parr *et al.*, 2014 | Velders *et al.*, 2012; Tchoukouegno Ngeu, 2013; Parr *et al.*, 2014; Chidi-Ogbolu & Baar, 2019 |
| **Fat-reducing / Hypolipidemic** | Santos *et al.*, 2012; Andrade *et al.*, 2014; Santos & Andrade, 2014; Schuchard *et al.*, 2015 | Singh *et al.*, 2012; Yadav *et al.*, 2013 | Syrov *et al.*, 1983; Kizelsztejn *et al.*, 2009; Seidlova-Wuttke *et al.*, 2010; Foucault *et al.*, 2012 | Seidlova-Wuttke *et al.*, 2010: Lizcano & Guzmán, 2014 |
| **Antidiabetic** | Liu *et al.*, 2012; Echeverria-Rodriguez *et al.*, 2014; Santos *et al.*, 2014; He *et al.*, 2015; Kilarkaje *et al.*, 2013 ; Chodavarapu & Lazartigues, 2015 ; Verma *et al.*, 2012 | Singh *et al.*, 2012 | Yoshida *et al.*, 1971; Kizelsztejn *et al.*, 2009; Sundaram *et al.*, 2012a,b | Mauvais-Jarvis, 2011 ; Zhang *et al.*, 2015; |
| **Anti-fibrotic** | Lubel *et al.*, 2009, Simões e Silva *et al.*, 2013; Barroso *et al.*, 2015; Willey *et al.*, 2016 | Phua *et al.*, 2009 | Hung *et al.*, 2012 | Wu *et al.*, 2009 |
| **Anti-inflammatory** | Da Silveira *et al.*, 2010; El-Hashim *et al.*, 2012; Simões e Silva *et al.*, 2013; Barroso *et al.*, 2015 ; Xue *et al.*, 2019 | Da Silvera *et al.*, 2010; Jawien *et al.*, 2012 ; Skiba *et al.*, 2017 | Kurmukov & Syrov, 1988; Xia *et al.*, 2016; Song *et al.*, 2019 | Pedersen *et al.*, 2016; Pelekanou *et al.*, 2016 |
| **Neuroprotective** | Jiang *et al.*, 2013; Regenhardt *et al.*, 2013; Zheng *et al.*, 2014; Bennion *et al.*, 2015; Villalobos *et al.*, 2016 | Lee *et al.*, 2015 ; Jiang *et al.*, 2018 ; Mo *et al.*, 2019 | Luo *et al.*, 2009; Liu *et al.*, 2011; Hu *et al.*, 2012 | Raz *et al.*, 2008; Lebesgue *et al.*, 2010 ; Arevalo *et al.*, 2015 |
| **Cardioprotective** | Tallant *et al.*, 2005; Benter *et al.*, 2007; Hao *et al.*, 2015 ; Tesanovic *et al.*, 2010. | Ferreira *et al.*, 2007; Ebermann *et al.*, 2008; He *et al.*, 2010; Zeng *et al.*, 2012; Cunha *et al.*, 2013; Yadav *et al.*, 2013; Ma *et al.*, 2016 | Kurmukov & Ermishina, 1991; Korkach *et al.*, 2007; Xia *et al.*, 2013a | Cong *et al.*, 2013, 2014 |
| **Vasorelaxant** | Dos Santos & Sampaio, 2015; Raffai *et al.*, 2011; Tesanovic *et al.*, 2010 | Lemos *et al.*, 2005 |  | Zhou *et al.*, 2013; Hermenegildo *et al.*, 2011 ; Sobrino *et al.*, 2010, 2017. |
| **Hematopoiesis stimulation** | Rodgers & di Zerega, 2013; Rodgers *et al.*, 2013 |  | Syrov *et al.*, 1997 | Nakada *et al.*, 2014 |
| **Liver protective** | Lubel *et al.*, 2009 ; Pereira *et al.*, 2007; Li, 2013 | Suski *et al.*, 2012 | Shakhmurova *et al.*, 2010a; Xia *et al.*, 2013b | Tian *et al.*, 2012 |
| **Lung protective** | Imai *et al.*, 2008; Klein *et al.*, 2013; Uhal *et al.*, 2013; Shenoy *et al.*, 2015 ; Cao *et al.*, 2019 | Klein *et al.*, 2013; Rodrigues-Machado *et al.*, 2013; Cao *et al.*, 2019 | Wu *et al.*, 1998; Li *et al.*, 2013; Xia *et al.*, 2016; Song *et al.*, 2019 | Hamidi *et al.*, 2011; Breithaupt-Faloppa *et al.*, 2013 |
| **Kidney protective** | Zhou *et al.*, 2012; Xu *et al.*, 2013; | Barroso *et al.*, 2012; Suski *et al.*, 2013 ; Silveira *et al.*, 2010, 2013 ; Pinheiro *et al.*, 2004. | Syrov *et al.*, 1992; Zou *et al.*, 2010; Hung *et al.*, 2012 | Iran-Nejad *et al.*, 2015; Wu *et al.*, 2016 |
| **Gastric protective** | Zhu *et al.*, 2014; Pawlik *et al.*, 2016 | Pawlik *et al.*, 2016 | Shakhmurova *et al.*, 2010b; Zhou *et al.*, 2010 | Du *et al.*, 2010 ; Liu *et al.*, 2010 |
| **Bone protective** | Krishnan *et al.*, 2013 |  | Gao *et al.*, 2008; Kapur *et al.*, 2010; Seidlova-Wuttke *et al.*, 2010b; Dai *et al.*, 2015 | Seidlova-Wuttke *et al.*, 2010b |

